# Actin and vimentin jointly control cell viscoelasticity and compression stiffening

**DOI:** 10.1101/2025.01.01.630993

**Authors:** James P. Conboy, Mathilde G. Lettinga, Pouyan E. Boukany, Fred C. MacKintosh, Gijsje H. Koenderink

## Abstract

The mechanical properties of cells are governed by the cytoskeleton, a dynamic network of actin filaments, intermediate filaments, and microtubules. Understanding the individual and collective mechanical contributions of these three different cytoskeletal elements is essential to elucidate how cells maintain mechanical integrity during deformation. Here we use a custom single-cell rheometer to identify the distinct contributions of actin and vimentin to the viscoelastic and nonlinear elastic response of cells to uniaxial compression. We used mouse embryonic fibroblasts (MEFs) isolated from wild type (WT) and vimentin knockout (vim -/-) mice in combination with chemical treatments to manipulate actin polymerization and contractility. We show through small amplitude oscillatory measurements and strain ramp tests that vimentin, often overlooked in cellular mechanics, plays a role comparable to actin in maintaining cell stiffness and resisting large compressive forces. However, actin appears to be more important than vimentin in determining cellular energy dissipation. Finally we show by comparing wild type and enucleated cells that compression stiffening originates from the actin and vimentin cytoskeleton, while the nucleus appears to play little role in this. Our findings provide insight into how cytoskeletal networks collectively determine the mechanical properties of cells, providing a basis to understand the role of the cytoskeleton in the ability of cells to resist external as well as internal forces.

**Significance statement:**

- A cell’s response to mechanical stress is largely governed by the actin and vimentin cytoskeletal networks, but their relative contribution to cell viscoelasticity and response to large deformations are poorly characterized.
- We reveal that actin and vimentin networks have an almost equal contribution to cellular stiffness and the cell’s ability to strain-stiffen under uniaxial compression.
- This work underscores the cytoskeleton’s central role in cellular mechanics and the mechanical synergy between the cytoskeletal networks, providing a framework for understanding how cellular components coordinate to maintain structural integrity and respond to different mechanical environments.

## Introduction

In multicellular organisms, cells are continuously exposed to different physical stresses. In some tissues and organs, cells are constantly subject to stretching (e.g. the heart, lungs and skin), while in others they are subject to compression (e.g. joints and the spine). ^1^ Cells can also actively generate forces to deform themselves, for instance during migration, which exposes cells to a complex combination of shear, tensile and compressive forces. ^2^ Furthermore, cells are able to stress-stiffen in response to mechanical stimuli. ^3^ In mammalian cells, the building blocks responsible for generating and responding to mechanical stress are the nucleus ^4^ and the cytoskeleton, a biopolymer scaffold comprised of three distinct components: actin, microtubules and intermediate filaments.^5^

In animal cells, filamentous actin is primarily located in the cell cortex, a dense meshwork that lines the plasma membrane, and in stress fibers, linear arrays of actin filaments that span the length of a cell. ^6^ In conjunction with myosin motor proteins, both the actin cortex and actin stress fibers are capable of generating contractile forces, termed cellular prestress, which has been shown to modulate cell stiffness. ^7^ Intermediate filaments are a large family of proteins expressed in a tissue-specific manner. The two most extensively researched intermediate filaments are keratin, which is found in non-motile epithelial cells, and vimentin, which is present in motile mesenchymal cells. ^8^ In mesenchymal cells, such as fibroblasts, vimentin intermediate filaments organize into a cage-like structure that envelops the nucleus and radiates outward towards the cell membrane. ^9^ Vimentin has been likened to a ’cellular safety belt’ that protects cells against large mechanical strains. ^10^ Microtubules also form a cell-spanning network, ^11^ but due to their limited abundance in the cell and their prevalence mainly as isolated filaments, they do not significantly contribute to whole-cell stiffness. ^12^

Understanding the relative contributions of actin and vimentin to cell rheology, as well as the potential mechanical synergy between these cytoskeletal networks, has long been a challenging question. Although actin and vimentin are widely recognized as key players in determining the cell’s mechanical response to external stress, precisely how each network contributes to cell elasticity and energy dissipation remains unclear. Previous studies have demonstrated that both actin^13^ and vimentin^14^ are integral to the material properties of cells, but their specific roles and interplay in these mechanical behaviors are yet to be fully understood.

Probing a cell’s mechanics at the nanometer to the micrometer scale has been performed via a plethora of techniques including optical tweezers, ^15,16^ atomic force microscopy,^17^ magnetic twisting cytometry^18^ and particle tracking microrheology.^19–21^ However, these localized techniques probe either mainly actin or mainly intermediate filaments, rather than their combined response. For example, mechanical measurements obtained from magnetic twisting cytometry are dominated by actin rather than vimentin because deformations are applied to the cell membrane, where the actin cortex provides the primary mechanical resistance. ^14,22^ Generally, due to the highly heterogeneous intracellular environment, it is difficult to extrapolate these microrheological results to a whole-cell mechanical response. Furthermore these local measurement techniques are often limited to small nanometer scale deformations, whereas cells in the body can experience rather large deformations. For instance, lung tissue cells are subjected to large cyclical forces from breathing. To access the large deformation regime, single-cell stretching and compression assays have previously been developed, where cells are either suspended between adhesive ^3,23^ or non-adhesive ^24^ parallel plates. However, these studies for the most part ignored the role of intermediate filaments in the cell’s mechanical response to large strain deformations. Intriguing recent work indicates that intermediate filaments provide a major contribution to the mechanical response of cells under large compressive strains. ^25^ This is consistent with observations from cell-free reconstitution experiments, which have shown that networks of intermediate filaments can carry much larger strains and exhibit more pronounced strain-stiffening than actin networks. ^26,27^ Here, we dissect the contributions of the actin and vimentin cytoskeletal networks to the nonlinear mechanical behavior of single fibroblasts by means of uniaxial compression rheology experiments on single fibroblasts confined between two nonadhesive parallel plates. To quantify the relative contributions of the two cytoskeletal networks, we used mouse embryonic fibroblasts isolated from wild type (WT) and vimentin knockout (vim -/-) mice in combination with drugs to influence actin polymerization (cytochalasin D) and contractility (blebbistatin). We show through small amplitude oscillatory measurements that the actin and vimentin networks equally contribute to total cell stiffness. However, actin plays a larger role than vimentin in determining cellular energy dissipation. Furthermore, we show that the strain-stiffening behaviour of cells arises from the actin and vimentin cytoskeleton rather than from the nucleus.

## Results

### Frequency-dependent rheology of single mouse embryonic fibroblasts

To quantify the macroscopic viscoelastic properties of single cells, we established a single cell rheometer assay for dynamic mechanical analysis on cells suspended between two parallel surfaces (Figure 1 A, top). Using an optical fiber with a wedge, we apply an oscillatory compressive strain of defined amplitude and frequency and measure the concomitant stress, which is used to calculate the frequency-dependent complex apparent Young’s modulus, *E*^∗^(*f* ) (Figure 1 A, bottom). We first validated the setup by benchmarking the system with polyacrylamide (PAA) microgels, which we used as a cell-sized proxy with well-characterized rheological properties^28^ (Figure 1 B, top). Our experiments showed that the elastic (storage) modulus, *E*^′^, of the microgels was constant between 0.1-10 Hz, and the viscous (loss) modulus, *E*^′′^, increased with the frequency of the applied strain (Figure 1 B, bottom), consistent with the predominantly elastic behavior of PAA as measured with bulk rheometry and atomic force microscopy.^28^

**Figure 1:**
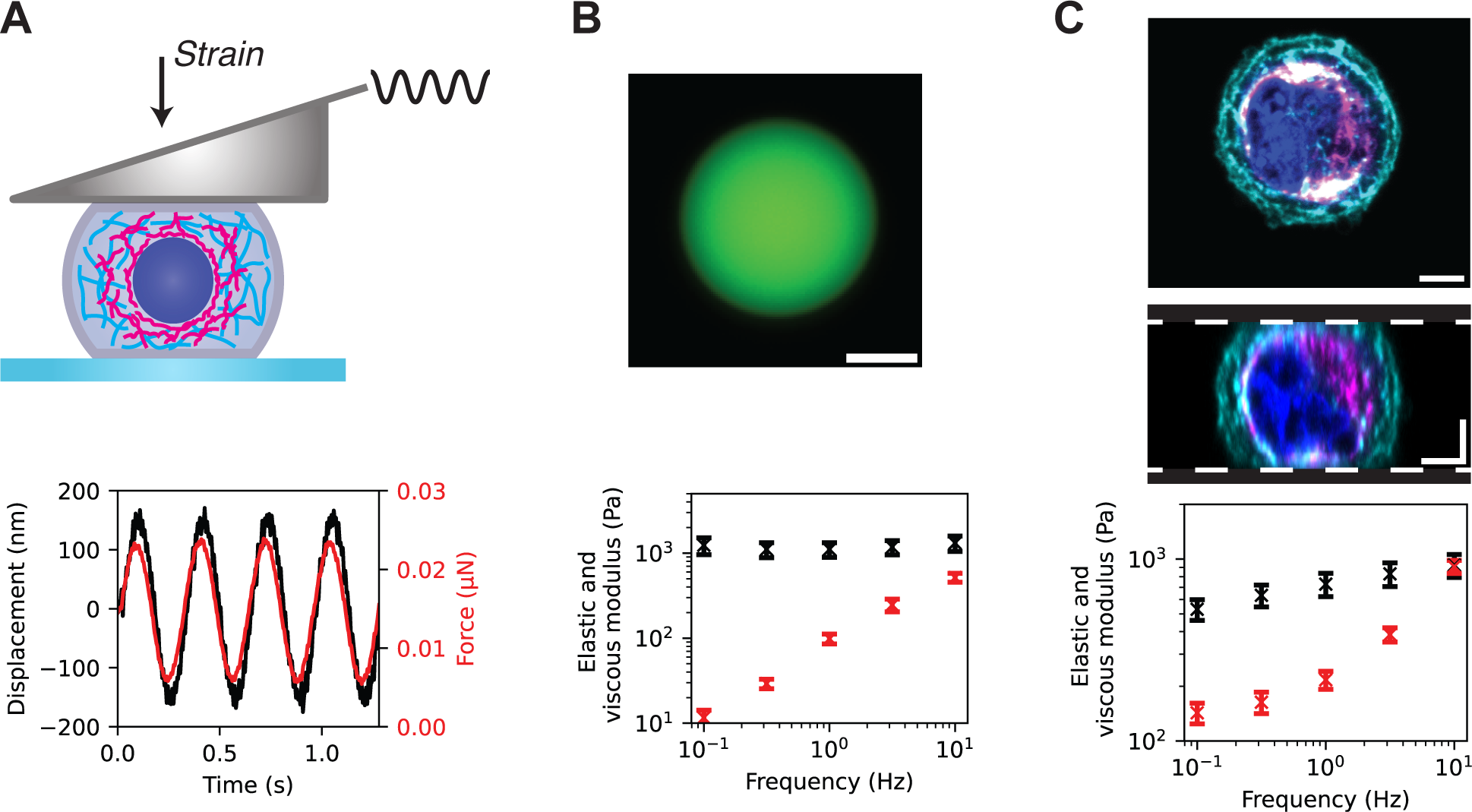
Oscillatory compression measurements of the linear viscoelastic properties of mouse embryonic fibroblasts (MEFs). (A) Top: schematic of a single MEF cell sandwiched between a glass coverslip and wedged flexible cantilever. We focus on the mechanical contributions of the actin cytoskeleton (cyan), vimentin cytoskeleton (magenta), and nucleus (dark blue). Bottom: representative measurement at an oscillation frequency of 0.32 Hz, showing the applied displacement of the cantilever (black; left y-axis) and observed force (red; right y-axis) as a function of time. (B) Top: epifluorescence image of a spherical polyacrylamide microgel used as a cell-sized analogue for benchmarking the setup. Scale bar: 20 µm. Bottom: Frequency-dependent elastic and viscous moduli (black and red symbols, respectively) of polyacrylamide microgels (average for N=8). (C) Single confocal slice through the equator (top) and orthogonal projection through the centre (middle) of a live MEF cell confined between two coverslips (indicated by white dashed lines) obtained by confocal fluorescence imaging. SiR-labeled actin (cyan) is largely located at the cortex while vimentin-GFP (magenta) forms a cage around the nucleus (dark blue). Bottom: frequency-dependent elastic and viscous moduli (black and red symbols, respectively) of MEF cells (average for N=16). Scale bars: 5µm.

We next performed similar measurements on mouse embryonic fibroblasts (MEFs). In contrast to the microgels, the cells behaved as viscoelastic solids with a power law dependence of *E*^′^ on frequency (Figure 1 C). At the highest frequency of 10 Hz, *E*^′^ and *E*^′′^ crossed, indicating that the cell transitioned to a viscoelastic liquid response. This type of power law dependence is a well-established phenomenological behavior of mammalian cells. ^29^

### Actin and vimentin both contribute to bulk cell viscoelasticity

To understand the contributions of actin and vimentin to cell mechanics, we first used confocal microscopy to examine the organization of the two networks in the wild type (WT) MEF cells. As shown in Figure 1C, the cells are quasi-spherical with a radially uniform distribution of actin and vimentin. We therefore generated ’average cells’ by averaging maximum intensity projections of 2 µm thick confocal sections centered at the cell equator, rescaling each cell to correct for differences in cell size (Fig. 2 top row). Filamentous actin is largely located at the cell periphery lining the plasma membrane as expected, but also extends to the interior of the cell. By contrast, vimentin is located in between actin and the nucleus. However, the averaged radial profile of actin and vimentin signals show that there is also a significant overlapping region where both actin and vimentin are localized.

**Figure 2:**
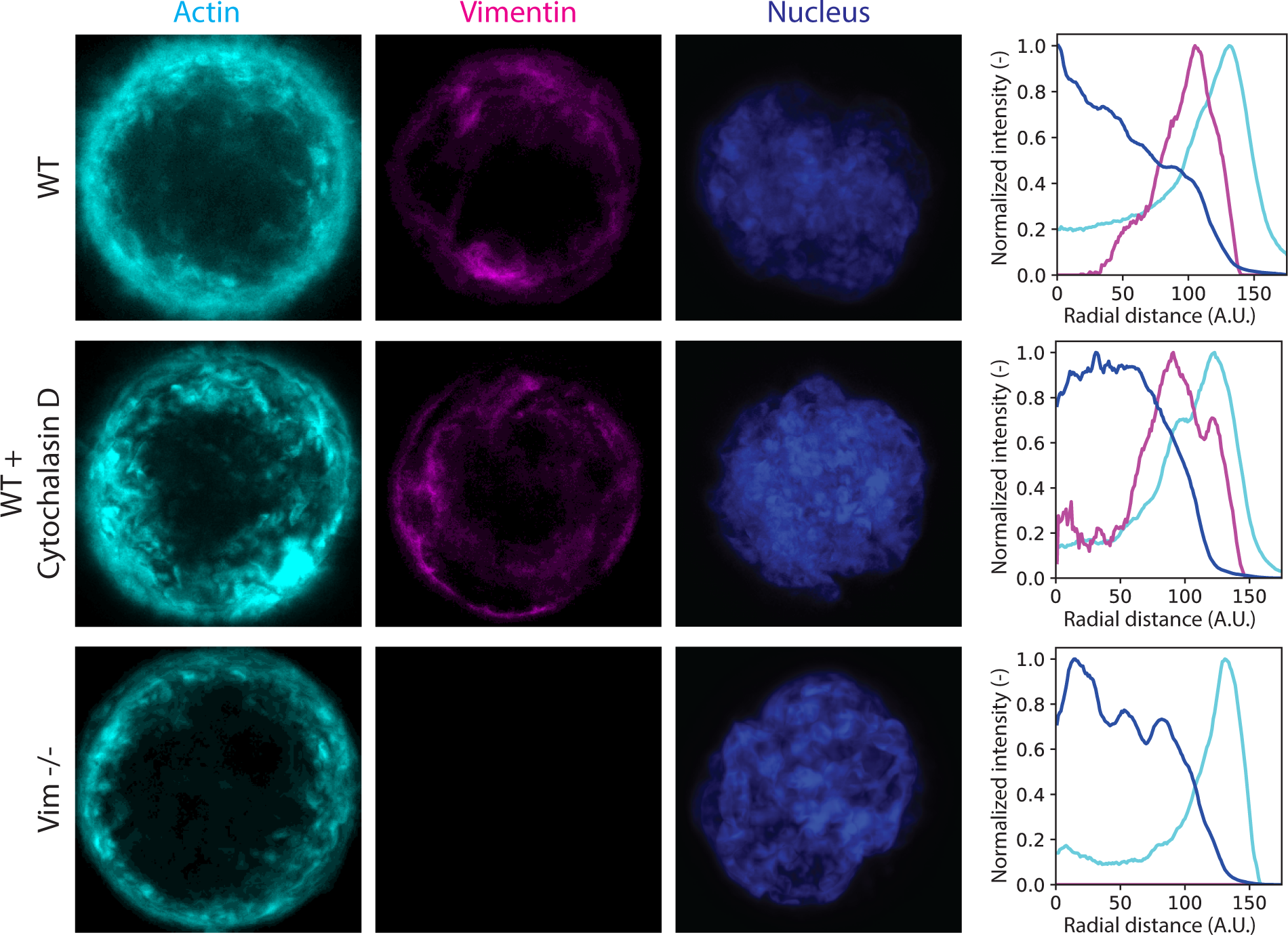
Actin (magenta), vimentin (cyan) and nuclear (dark blue) organization in ‘average’ WT MEF cells (top), WT cells drugged with cytochalasin D (middle), and vim -/cells (bottom). As we rescale cells to generate ‘average’ cells, scale bar not applicable. The images represent averages of a 2 µm thick confocal section at the cell equator for 10 cells. For all conditions, filamentous actin is located largely at the cell periphery and vimentin is located closer to the center of the cell. Average radial profiles of the fluorescent signals (right) were calculated by taking normalized intensities around concentric circles from the center of the image, which show a significant overlapping region between actin and vimentin. The radial distance is set to zero at the cell center; the distance is in arbitrary units (see Methods for details).

Next, to isolate the individual contributions of actin and vimentin to the mechanical response of the MEF cells, we set up experiments where we removed either F-actin or vimentin. To remove F-actin, we treated the WT cells with cytochalasin D.^30^ As shown in the averaged confocal images and radial intensity profiles in Figure 2 (middle row), there is a remaining filamentous actin signal after cytochalasin D treatment, but this signal is diffuse and less strongly located at the cortex than in WT cells. We do not observe any striking change in the localization of the vimentin network upon cytochalasin D treatment. To remove the contribution of vimentin, we turned to vim -/MEFs that are genetically depleted of vimentin. ^31^ As shown in the images and radial intensity profiles in Figure 2 (bottom row), the cells are indeed completely devoid of vimentin while the actin network appears somewhat more confined close to the membrane compared with WT cells.

To quantify the relative contributions of actin and vimentin to single cell viscoelasticity, we compared compression measurements on the wild type cells, cytochalasin D treated cells, and the vim -/cells. As shown in Fig. 3A, the viscoelastic moduli significantly decreased both upon drug-induced depolymerization of actin and upon knocking out vimentin. In order to make a quantitative comparison, we chose to model the complex compression modulus with the structural damping law, a well-known phenomenological model of cell rheology. ^32–34^ To separate out the contribution of the cytoplasm, we included an additive Newtonian viscous damping term:

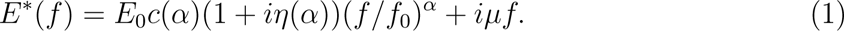

**Figure 3:**
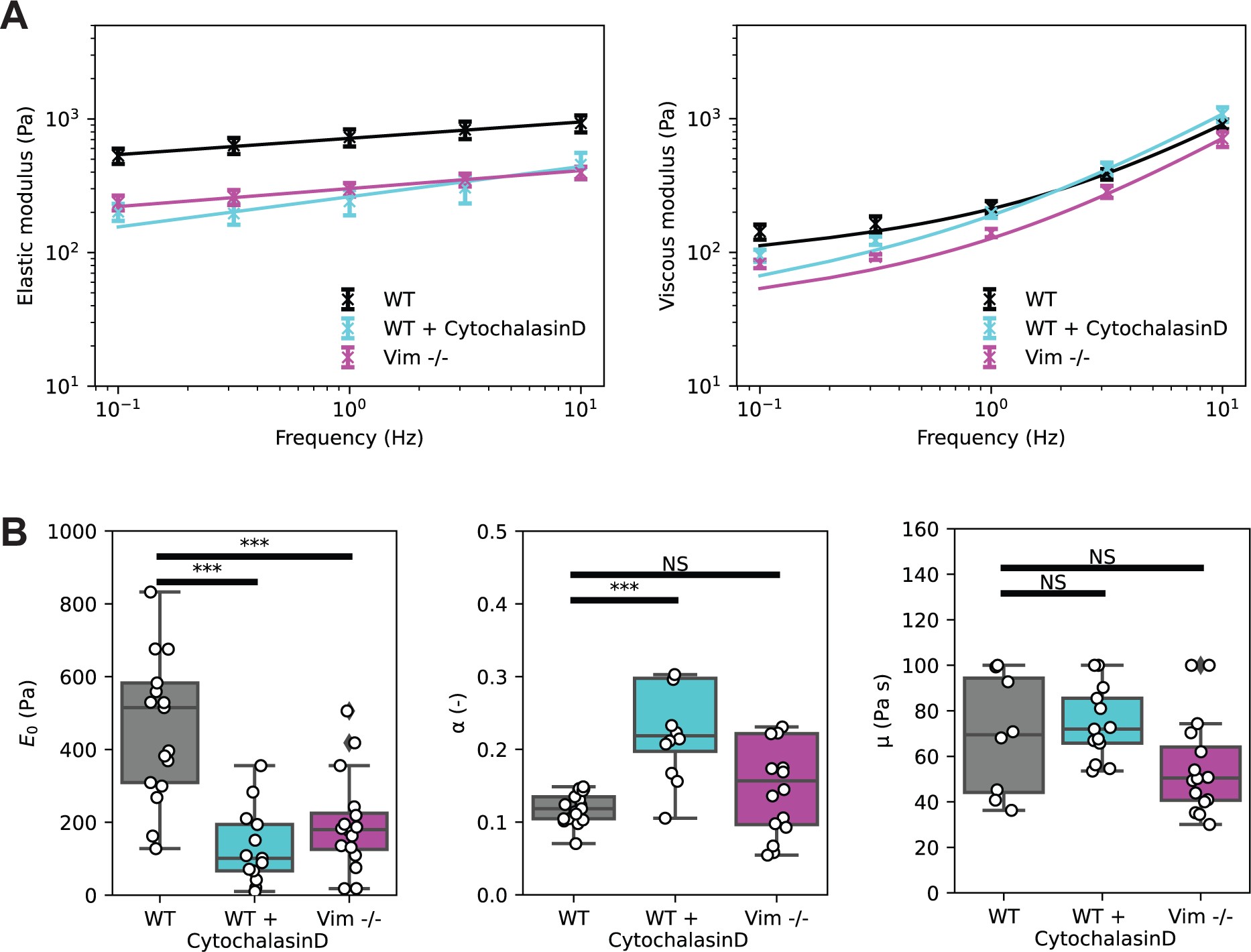
(A) Elastic (left) and viscous (right) linear compressive moduli (symbols), measured for oscillation frequencies between 0.1Hz and 10Hz, with fits (solid lines) to the structural damping model (Eq. 1). (B) The prefactor *E*_0_ from the viscoelastic fit shows a significant decrease, both after drug-induced depolymerization of actin and upon knocking out vimentin, as compared to wild type cells. (E) The power law exponent *α* increases only upon actin depolymerization. (F) Neither actin depolymerization nor vimentin knockout changes the Newtonian viscous damping term *µ*. **P<0.01, ***P<0.001. NS, not significant.

Here, *E*_0_ is a scaling factor for the elastic and viscous moduli at a frequency scale factor *f*_0_ (which we arbitrarily set to 1 Hz), *c*(*α*) is Γ(1 − *α*) cos(*πα/*2) where Γ denotes the gamma function, *η* = tan(*πα/*2) is the structural damping coefficient, *α* is the power law exponent and *µ* is the Newtonian viscous term. The great advantage of this model is its ability to reduce the frequency-dependent measurements to three parameters (*E*_0_, *α* and *µ*), which provide a mechanical fingerprint to allow comparisons between different conditions. Importantly, the structural damping law provides a self-consistent fit of *E*^′^ and *E*^′′^ to the real and imaginary parts of the complex modulus (lines in Figure 3A).

We find that the scaling factor *E*_0_ decreases from 505 ± 70 Pa for wild type cells to *E*_0_ = 130 ± 30 Pa for cells drugged with cytochalasin D and *E*_0_ = 196 ± 30 Pa for vimentin -/cells (see Fig. 3B left). This finding demonstrates that both actin and vimentin contribute to the bulk elasticity of the cell. Interestingly, if one were to model the *E*_0_ value for WT cells as the sum of two independent elastic contributions from the actin and vimentin cytoskeletal networks by adding the *E*_0_ values we measured for the cytochalasin-D treated cells and the vimentin -/- cells, one obtains a value (326 Pa) that is significantly less than the value measured for wild type cells. This suggests a mechanical crosstalk between the two networks. At the same time, the power law exponent *α* increased from *α* = 0.112 ± 0.005 for wild type cells to *α* = 0.30 ± 0.06 for actin-depolymerized cells. However, for vimentin -/- cells, we observed no significant change in *α* value (0.20 ± 0.04) as compared to wild type cells (see Fig. 3B middle). The measured exponents reveal viscoelastic behavior, being in between the limits of an ideal solid (*α* = 0) and an ideal fluid (*α* = 1). Depolymerization of actin causes the cells to become more fluid-like, indicating that actin is more important than vimentin in controlling energy dissipation. Finally, the Newtonian viscous damping term *µ* remained invariant for all cell perturbations (Fig. 3B right).

### Strain ramp experiments to characterize the nonlinear elasticity of MEF cells

Next, we sought to determine whether the stiffness of the cells changes with increasing compression. We applied a uniaxial compression where we steadily increased the strain from 10 zero to ∼50% at a fixed deformation rate of 1 µm/s (Figure 4A). We selected this rate to minimize potential viscous effects and ensure that the cell’s mechanical response remained primarily elastic. We limited the compressive strain to 50% because we observed that cells could tolerate repeated straining up to this point without any visible damage or changes in mechanics (Supplementary Video 1). By contrast, cells often forms blebs indicative of loss of viability at strains when we compressed them beyond this point (see Supplementary Video 2).

**Figure 4:**
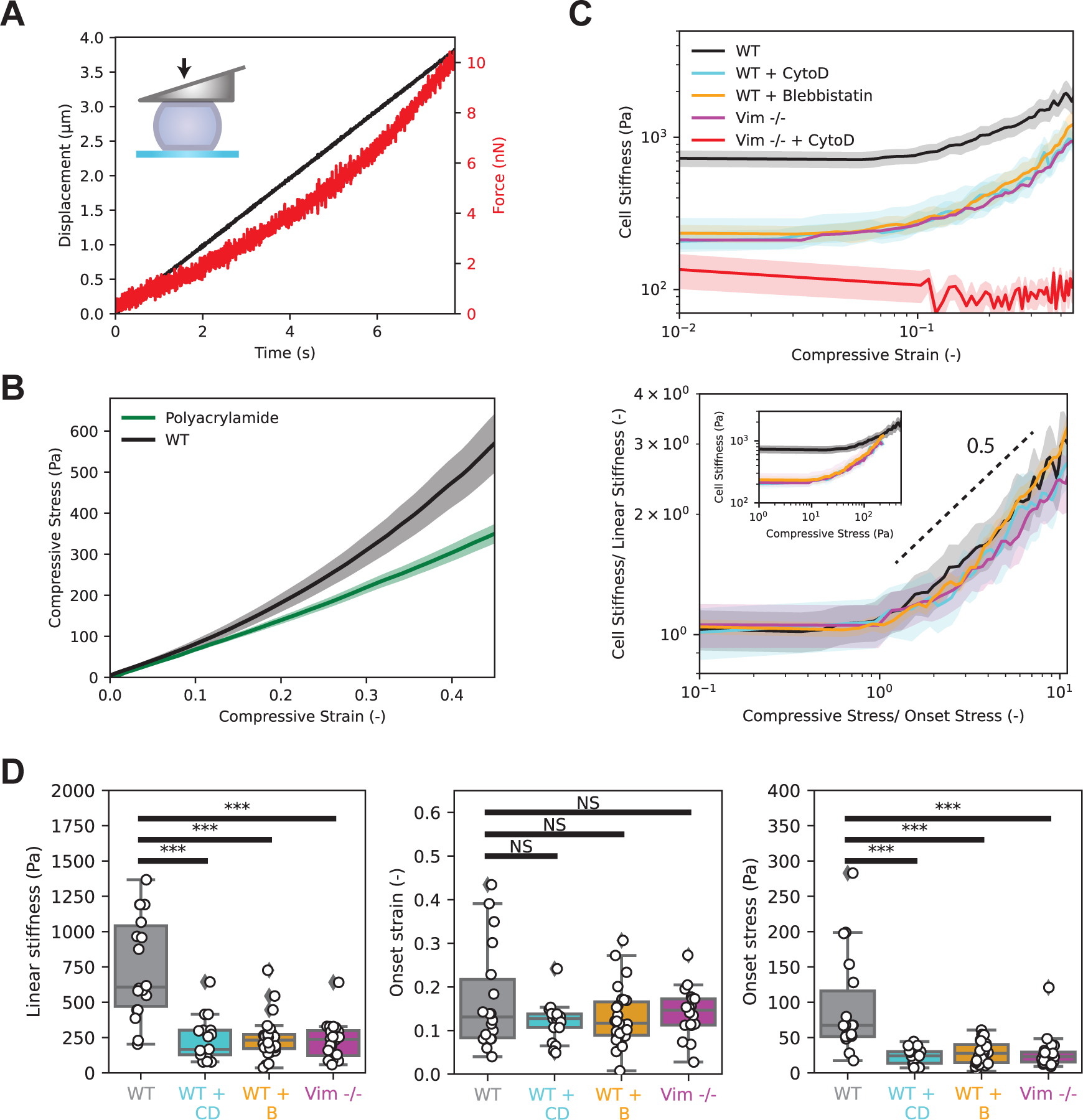
Strain ramp experiments to characterize the nonlinear response of MEFs to large compressions. (A) Representative strain ramp measurement showing the applied displacement of the cantilever (black, left y-axis) and observed force (red, right y-axis). (B) Applied strain versus measured stress for wild type cells (N=18) and polyacrylamide microgels (N=8). Cells strain-stiffen whereas polyacrylamide gels show a linear response. Shaded areas around the curves show the standard error. (C) Top: Cell stiffness versus strain, obtained by taking the numerical derivative of the stress-strain curves from panel B. Wild type cells (untreated (N=18) or treated with either blebbistatin (N=26) or cytochalasin D (N = 17) and vimentin KO (N=20) cells strain-stiffen. Vimentin KO cells treated with cytochalasin D (N=14) are significantly softer and no longer strain-stiffen. Bottom: Normalizing the stress by the onset stress for stiffening and the cell stiffness by the linear stiffness, respectively, collapses the curves onto a single master curve. The dashed line indicates power-law stiffening with an exponent of 0.5. Inset shows the same data without rescaling. (D) Left: All drug and genetic perturbations lead to a significant decrease in the linear stiffness of the cells. Middle: The onset strain for strain-stiffening is invariant. Right: all drug and genetic perturbations lead to an earlier onset stress for strain-stiffening. Shaded areas around each line show the standard error on the mean. ***P<0.001. NS, not significant.

We first benchmarked the assay by compressing the same microgel beads also used for benchmarking the oscillatory measurements. We found that the stress increased linearly with strain (green curve in Figure 4B), equivalent to the linear elastic response of polyacrylamide as measured in bulk rheometry.^35^ In stark contrast, the cells showed a marked strain-stiffening response (black curve in Figure 4B). To characterize this response, we calculated the nonlinear stiffness *K* of the cells to compressive loading in a way similar to what has been done for shear stress in reconstituted networks, as a derivative of the stress with respect to strain, rather than the ratio of stress to strain in the case of the linear modulus. ^29,36^ Specifically, 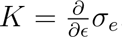, where *ɛ* is the compressive strain and *σ_e_* is the resulting compressive stress measured in Figure 4B. From this procedure, we found that wild type MEF cells exhibited a linear regime for compressive strains below ∼10%, followed by nonlinear stiffening (Figure 4C). This finding shows that cells compression-stiffen under uniaxial load, consistent with prior reports.^37^

### Actin and vimentin both contribute to the compression-stiffening of cells

To test the contribution of the actin cytoskeletal network to the nonlinear stiffening behaviour of the cells, we then performed strain ramp measurements on wild type cells drugged with cytochalasin D. Actin depolymerization resulted in a marked softening of the cells over the entire strain range compared to wild type cells, but the cells continued to exhibit strain-stiffening behaviour (cyan curve in Figure 4C). The linear modulus dropped from 747 ± 80 to 230 ± 30 Pa, the onset strain for strain-stiffening did not change significantly, and the onset stress was reduced (Figure 4D). When we performed the same experiment on vimentin knockout cells, we again observed a lower linear elastic modulus (234 ± 40 Pa) compared to wild type cells, invariant onset strain and a lower onset stress (magenta curve in Figure 4C, summary in Figure 4D). Our findings indicate that actin and vimentin both contribute to the strain-stiffening behavior of MEF cells. To test whether strain-stiffening is preserved when both networks are removed, we measured the response of vimentin knockout cells drugged with cytochalasin D to strain ramps. Strikingly, we observed a complete loss of strain stiffening behaviour in addition to a strong drop of the elastic modulus (red curve in Figure 4C). We note that, despite this drastic treatment to the cytoskeleton, cell viability was not impacted (Figure S3). Therefore, we conclude that both the actin and vimentin networks are responsible for the compression stiffening of the cells.

It is known that the stiffness of mammalian cells is strongly influenced by the level of internal contractile stress *σ_p_* (also known as prestress) generated by nonmuscle myosin-2 motor activity.^29^ It has also been shown that motor-generated stress and externally applied stress *σ_e_* play similar roles in the stiffening of reconstituted actomyosin networks, ^38^ as has also been shown in simulations of model fiber networks. ^39^ It has further been suggested that the nonlinear increase in the modulus is proportional to the sum of internal and external stress *σ_p_* + *σ_e_*. According to this phenomenological model, the nonlinear stiffness would increase linearly with applied stress. To investigate the dependence of cell stiffness (in Fig. 4C) on applied stress, we rescale *K* by the linear value (i.e., in the limit of small stress and strain) and plot this by the ratio of the stress to its value at the onset of stiffening. Importantly, this rescaling procedure is found to result in a good collapse of all data to a single master curve that characterizes both wild type cells and cells with different drug or genetic manipulations.

In the stiffening response to stress, we observe an apparent power-law stiffening exponent of ∼ 0.5 (Figure 4C bottom). While this differs from the linear dependence suggested in prior work, we note that such an exponent has been predicted theoretically for fibrous networks^39–41^ and seen experimentally for other biopolymer networks. ^42^ The onset stress is reduced for both actin-depleted and vimentin knock out cells (Figure 4D). Indeed, assuming a given onset tension per filament, the removal of network filaments is expected to lead to both a reduction in the linear modulus and in the onset stress. In the case of actin, depletion can also reduce the prestress *σ_p_*. In order to identify the role of prestress, we also treated wild type cells with blebbistatin, which inhibits myosin-2. As shown in Fig. 4 (yellow curve), the blebbistatin-treated cells showed an identical strain-stiffening response to cells treated with cytochalasin D (cyan curve), indicating that myosin-generated prestress dominates the mechanical contribution of actin. Thus, the higher onset stress of wild type cells can be accounted for by their higher contractile prestress *σ_p_*.

### The cytoskeleton, not the nucleus, causes compression stiffening

From the imaging experiments, we observed that the nucleus occupied a large volume fraction (∼60%) of our cells (Figure 2). While the nucleus has long been regarded as the stiffest component of the cell, ^43^ recent studies suggest that it may actually be softer than the cell as a whole. ^44–46^ Therefore, we sought to distinguish the response of the cytoskeleton from any stiffening or softening effects that could be caused by compression of the nucleus. To this end, we removed the nucleus from wild type MEF cells by centrifuging cells plated on coverslips following a well-established procedure. ^47^

As shown in Figure 5A, the resulting enucleated cells (also known as cytoplasts) retain an actin and vimentin cytoskeleton with similar architecture to that of WT cells. When we performed strain ramps on the enucleated cells, we found that they still strain-stiffened, although they were softer than the intact cells (Figure 5B). The stress-stiffening curves of intact and enucleated cells collapsed onto a single master curve upon normalizing the stress and modulus axes (Figure 5B inset), showing that the stiffening is controlled by the cytoskeleton. We conclude that the cells stiffen under macroscopic compression due to the nonlinear response of the cytoskeleton rather than as a consequence of the nucleus.

**Figure 5:**
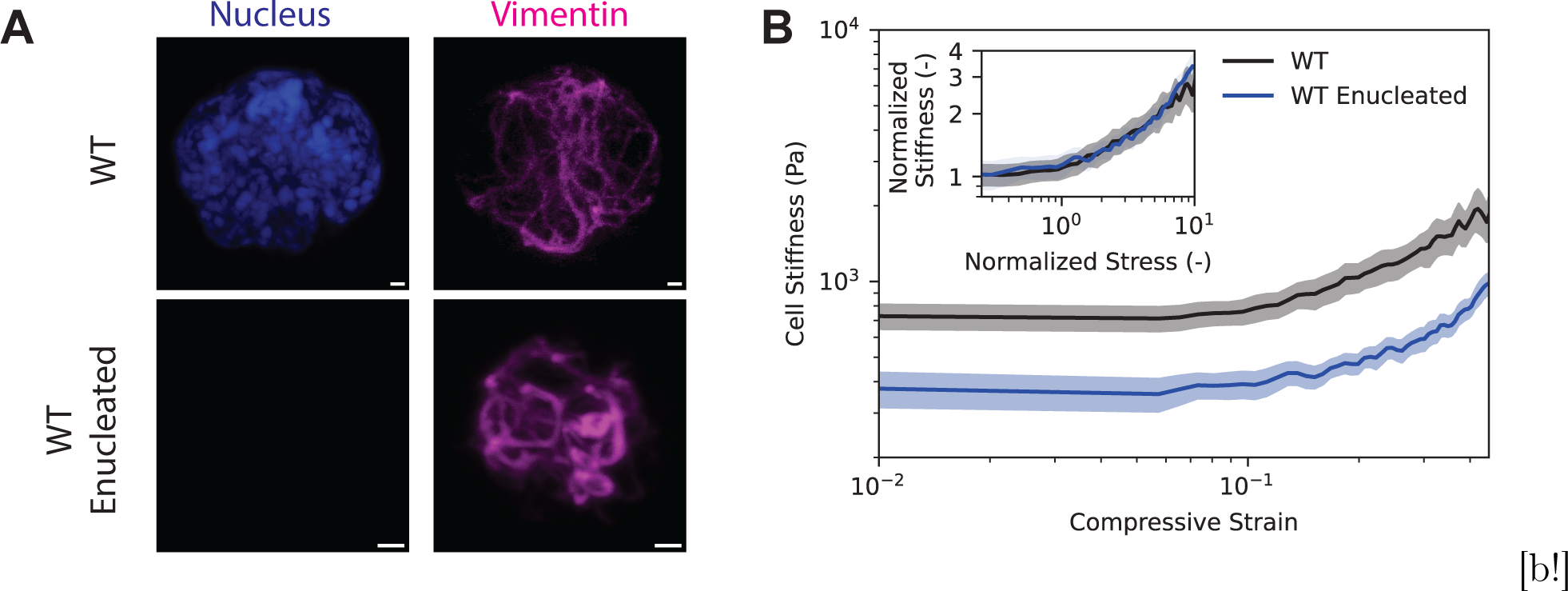
Strain ramp experiments on intact and enucleated wild type cells. (A) Maximum projection confocal images of the nucleus and vimentin in wild type (top) and enucleated wild type (bottom) cells. Enucleated cells no longer have a nucleus but vimentin still is able to form a network. (B) Wild type (N=18) and enucleated cells (N=15) both strainstiffen, with the same onset strain for stiffening, but enucleated cells are softer. Inset: The stress-stiffening curves collapse onto a single master curve upon normalization by the linear stiffness and by the onset stress for stress-stiffening. Shaded areas around the curves show the standard error on the mean.

## Discussion

In this study, our aim was to quantify the viscoelastic properties and nonlinear elasticity of single mouse embryonic fibroblasts (MEFs) and to dissect the contributions of the actin and vimentin cytoskeletal networks. Using a custom single-cell rheometer, we measured the linear viscoelastic properties through small amplitude oscillatory compression, where we controlled the applied strain and measured the concomitant stress. Using polyacrylamide (PAA) microgels, we validated the rheometer setup, showing expected elastic and viscous responses consistent with bulk measurements and linear elastic behavior. Our oscillatory measurements revealed that single fibroblasts behave as viscoelastic solids, with a frequency-dependent elastic modulus, *E*^′^, that increases with frequency following a power-law, a characteristic feature of mammalian cells. Beyond small deformations, we applied large uniaxial strains to assess nonlinear cell elasticity. In these measurements, we found that the cells exhibit pronounced strain-stiffening, with the differential elastic modulus increasing steeply beyond 10% compressive strain. Both small and large strain experiments highlighted the critical roles of actin and vimentin in cell mechanics. Depolymerizing actin or knocking out vimentin led to significant reductions in both the linear viscoelastic moduli and the nonlinear strain-stiffening response, underscoring that both networks contribute to the cell’s resistance to deformation, especially under large mechanical loads.

Our mechanical fingerprinting allowed us to distinguish the contributions of actin and vimentin to cell viscoelasticity. Our small amplitude oscillatory measurements show that the WT MEFs are 3-fold stiffer as compared to cells lacking actin (WT MEFs drugged with cytochalasin) or vimentin (Vim -/- MEFs). Although actin has traditionally been considered the dominant contributor to cell stiffness, we found that vimentin and actin make comparable contributions to the linear and nonlinear elasticity of suspended MEF cells. Our measured cell stiffness values (*E*_0_) are about one order of magnitude lower than values reported for cells adhered to glass, ^48^ but comparable to prior measurements on fibroblasts in suspension ^49^

(see Table 1). The discrepancy with adherent cells is likely due to the lack of visible actin stress fibers in suspended cells. In contrast, adherent MEF cells form pronounced stress fibers (see Supplementary Figure 2), , which are known to contribute to stiffness in adhered cells. ^50^ Previous studies showed that the measurement conditions also significantly affect cell stiffness. For instance, Fernández et al.^3^ reported stiffness values of 15 kPa for 3T3 fibroblasts using microplate rheology, whereas Mendez et al.^48^ measured smaller values of 3.5 kPa for wild type and 2.9 kPa for Vim -/- fibroblasts using atomic force microscopy. In both studies, the cells were adherent to a rigid substrate, which increases cell stiffness. Fibroblasts in suspension generally exhibit much lower stiffness. For example, Gerum et al.^49^ reported 85 Pa for WT cells, decreasing to 75 Pa with cytochalasin D treatment, and even lower values (60 Pa without and 35 Pa with cytochalasin D) for Vim -/- cells. These values align with our findings, suggesting that suspended fibroblasts, regardless of cytoskeletal changes, are softer than adherent fibroblasts. Differences between our study and Gerum et al.’s could stem from variations in MEF cell lines and the use of different measurement methods. While we employed uniaxial compression, Gerum et al. used shear-based flow cytometry.

**Table 1:** Summary of previously measured stiffness values of mouse embryonic fibroblasts.

| Reference | Cell Stiffness | Method |
| --- | --- | --- |
| Fernández et al.<sup>3</sup> | 15 kPa 3T3 and 3 kPa 3T3 + Latrunculin A | Microplate rheology<sup>†</sup> |
| Mendez et al.<sup>48*</sup> | 3.5 kPa WT and 2.9 kPa Vim -/- | Atomic force microscopy<sup>†</sup> |
| Guo et al.<sup>14*</sup> | 10 Pa WT and 5 Pa Vim -/- | Optical tweezers<sup>†</sup> |
| Gerum et al.<sup>49</sup> | 85 Pa 3T3 and 75 Pa 3T3 + CytoD<br>60 Pa Vim -/- and 35 Pa Vim -/- + CytoD | Flow cytometry<sup>‡</sup> |

<sup>\*</sup>These are the same cells used in this study. <sup>†</sup>Cells in these studies were measured in an adherent state. <sup>‡</sup>Cells in these studies were measured in an suspended state.

Interestingly we found that vimentin knockout altered the elastic component of the complex modulus but did not significantly change the power law scaling exponent *α* describing the frequency dependence. By contrast, actin depolymerization with cytochalasin D did change *α*. This observation indicates that actin plays a larger role than vimentin in controlling cellular energy dissipation. This is consistent with atomic force microscopy (AFM) experiments performed on bone-marrow-derived dendritic cells, where no significant difference was found in stress relaxation times between wild-type and vimentin knockout cells. ^51^ It is also consistent with bulk rheology measurements on reconstituted actin-vimentin composite networks, where vimentin was shown to be less dissipative than actin.^52^ These findings also align with the known difference in stability of actin and vimentin networks in the cell. Actin networks are highly dynamic, with a typical turnover time on the order of minutes in case of stress fibers ^53^ as well as the actin cortex.^54^ By contrast, intermediate filament networks exhibit turnover times on the order of hours. ^55^ This picture is consistent with previous studies showing that vimentin intermediate filaments form a stretchable, hyperelastic network that enhances cytoplasmic resilience and toughness under mechanical stress. ^56^

Our strain ramps combined with cytoskeleton perturbations and enucleation experiments point toward the cytoskeleton as the main component responsible for strain stiffening, with comparable contributions from actin and vimentin. Both purified actin and vimentin networks have indeed been shown to exhibit strain-stiffening as a consequence of the semiflexible polymer nature of the filaments. Recent evidence suggests that vimentin plays a dominant role in determining the stiffening behaviour of cells, ^25^ but we observed that both actin and vimentin contribute to this stiffening behaviour. In the stiffening response to stress in Figure 4C bottom, we observe an apparent power-law stiffening exponent of ∼ 0.5. Such an exponent has been predicted theoretically for networks^39–41^ and seen experimentally for other biopolymer networks. ^42^ This behavior can be understood in simple terms as follows. If the stiffening is due to additional network elements becoming mechanically engaged only above a critical threshold strain *ɛ_c_*, it is natural to expect that the stiffness *K* may increase approximately linearly in the excess strain *ɛ* − *ɛ_c_* above threshold. This would correspond to the stress contribution *σ* that would increase as (*ɛ* − *ɛ_c_*)^2^, leading to *K* ∼ *σ*^1^*^/^*^2^. While this behavior has previously been discussed in the context of shear strain and bulk expansion, it is important to note that cells are expected to maintain their volume under compression, which requires shear. Whether this mechanism is the origin of the approximate *K* ∼ *σ*^1^*^/^*^2^ scaling we observe is not yet clear. It may also be that a larger exponent closer to 1 may appear at higher stress. For cells under compression, however, the corresponding high strains may not be achievable without killing the cells.

Our compression experiments on enucleated cells show that the nucleus has a positive contribution to the elastic response of the cell to deformation. This is consistent with reports showing nuclei are stiffer than the surrounding cytoplasm. ^57^ However, our results show that the nucleus does not mediate the stress stiffening of single cells. Previous attempts to explain the rheology of cells in suspension have utilized a shell model approach. ^24^ The shell model in cellular rheology assumes that mechanical behavior is primarily dominated by the cell cortex, with the bulk of the cell’s mechanical response attributed to the elastic and viscous properties of this outer shell, rather than contributions from deeper cytoplasmic elements. However, our findings reveal that both vimentin and the nucleus significantly contribute to cell elasticity. Since these structural elements are located within the cell’s interior, this suggests that the cortex alone does not fully account for cellular mechanical behavior, indicating a more complex interplay of intracellular components in determining cell mechanics.

Our strain ramp experiments demonstrate that both actin and vimentin are key mediators of prestress. Cells are able to generate contractile stresses through the interaction of actin filaments and myosin motors, which gives rise to cellular prestress. ^58^ The strain ramp experiments suggest that actin depolymerization reduces the cellular prestress, which makes sense since contractility requires an intact actin filament network to allow force transmission. Interestingly, vimentin knockout also appears to reduce the cellular prestress. The reason for this effect is less clear. Clues for this have emerged from growing evidence that vimentin is a key regulator of actin organization in the cell cortex^59^ and in stress fibres. ^60^ Furthermore, previous traction force microscopy measurements have shown that wild type MEFs are ∼46% more contractile than their vimentin knockout counterparts. ^61^ Although these measurements were performed on adherent cells spread on flat substrates, in contrast to our suspended cell setup, our finding that vimentin knockout reduces cellular prestress is nevertheless consistent with the observation of reduced traction force. A loss of contractile ability upon vimentin depletion may help explain why cells without vimentin exhibit slower migration speeds as compared to WT cells. ^51^ Our findings are particularly relevant for understanding the ability of cells to migrate through narrow extracellular spaces, which requires deformation of both the nucleus and the cytoskeleton. ^62^ Understanding how these internal components modulate cell mechanics could lead to more accurate models of cancer cell migration and help develop targeted therapies that alter intracellular structures to inhibit metastatic spread.

Strikingly, in all of our experiments we found that total cell stiffness cannot be solely attributed to the individual contributions of actin and vimentin, suggesting a mechanical synergy between these filaments. A possible candidate for this synergy is crosslinking, likely mediated by the cytolinker plectin that forms crosslinks between actin and vimentin filaments.^63^ Crosslinkers generally increase the elastic modulus of cytoskeletal networks while reducing viscous dissipation. ^64^ Our recent rheology experiments of reconstituted crosslinked actin-vimentin networks, facilitated by a plectin-like cytolinker, revealed clear evidence of mechanical synergy between the filaments.^52^

sectionConclusion We demonstrated that actin and vimentin networks play complementary roles in determining the mechanical properties of mouse embryonic fibroblasts. Using a custom single-cell rheometer, we quantified the viscoelastic and nonlinear elasticity of suspended cells, revealing significant strain-stiffening behavior. Both actin and vimentin contribute to cellular stiffness, with vimentin also influencing prestress and hyperelasticity. Our results highlight the mechanical synergy between these two cytoskeletal networks and provide insight into how cytoskeletal organization impacts cell mechanics. This study offers a deeper understanding of the mechanical behavior of fibroblasts and emphasizes the importance of both actin and vimentin in cellular function and response to mechanical forces.

## Methods

### Cell culture

Wild type (WT) and vimentin-null (Vim -/-) mouse embryonic fibroblast (MEF) cells, obtained from WT and Vim -/- embryos, ^65^ were cultured in Dulbecco’s modified Eagle’s medium with glutamax (DMEM, 10565018 Gibco) supplemented with 10% fetal bovine serum (FBS, 10270106 Gibco) and 5% penicillin-streptomycin (Pen-Step, 15070063 Gibco). Cells were cultured in a humidified incubator maintained at 37°C and 5% CO_2_, and subcultured when they reached approximately 80% confluency. Cells were subcultured twice per week and were tested for mycoplasma contamination every four months. The day prior to mechanical loading experiments, 10000 cells, counted with a Countess cell counter (Thermo Fisher Scientific), were seeded in 6 plastic cell culture well plates (83.3920.005 Sarstedt) in standard cell culture medium. On the day of experiments, cells were detached with Trypsin-EDTA (0.25%) (25200056 Thermo Fisher Scientific) and resuspended in CO_2_-independent medium (18045088 Gibco) supplemented with 10% fetal bovine serum. For actin depolymerization experiments, CO_2_-independent medium was supplemented with Cytochalasin D (C8273 Sigma) at a concentration of 1 µM, chosen based on previous reports.^49^ For myosin inhibiting experiments, CO_2_-independent medium was supplemented with blebbistatin (203391 Sigma) at a concentration of 20 µM, also based on literature reports.^66^ Cells were incubated in drug-supplemented medium for at least 15 minutes before experiments.

### Fabrication of wedged force cantilevers

Tipless microcantilevers with a nominal spring constant of 0.018 N/m (Optics11 Life) were modified to correct for a 3^◦^ cantilever tilt^67^ and thus facilitate uniaxial compression. We first prepared coverslips, functionalized to prevent cell adhesion. 18 mm glass coverslips were cleaned by sonicating them in 2-Propanol (34863 Sigma) for 5 minutes and were then air-dried. The cleaned coverslips were next submerged in a solution of 10% v/v SurfaSil (TS42800 Thermo Fisher Scientific) in chloroform for 10 seconds. The coated coverslips were then air dried overnight on parafilm-coated 10cm petri dishes. We checked the efficacy of the Surfasil coating by placing a small drop of UV NOA81 curing polymer adhesive (NOA81 Norland) on the coverslip followed by curing using a UV (395 nm) LED (11504255 Leica) at a distance of approximately 5 cm. If the drop could be moved with a pipette tip without any difficulty, the coverslip could be used for cantilever fabrication. Next a small amount of NOA81 polymer adhesive was deposited on the microcantilever using the tip of a needle under a stereoscopic microscope. The cantilever was pressed against a coated coverslip to create a flat surface and the polymer adhesive was cured using the UV LED at a distance of approximately 5 cm for two hours. The resulting wedged cantilever was carefully removed from the coverslip by first gently tapping the microscope stage to promote detachment and then gradually moving the cantilever up in steps of 2 µm. Cantilevers were calibrated by acquiring a force-distance curve on a stiff surface (glass), according to the manufacturer’s instructions. This was performed before and after wedge manufacture to ensure no change in spring constant. After fabrication, the cantilever was cleaned with 2-Propanol, rinsed with MilliQ water and carefully dried with a Kimtech tissue.

### Single cell rheology experiments

The uniaxial compression setup consisted of an optical fiber-based nanoindenter (Optics11 Life) mounted on a Leica Thunder Imager widefield microscope equipped with a 200 mW solid state LED5 light source (Leica) and a monochrome sCMOS camera (Leica). For imaging, we used a 10x dry objective (HC PL APO 0.45NA, Leica). At the beginning of every measurement session, cantilevers were calibrated using the stiff-surface contact method, performed according to the manufacturer’s instructions. To prevent adhesion between cells and the culture dish, dishes were first cleaned with isopropanol, exposed to deep UV via a ProCleaner UV ozone cleaner (Bioforce Nanosciences), incubated for 1 hour in 0.1 mg/mL PLL-g-PEG solution (SuSoS), and finally washed with MilliQ water and allowed to air dry. The dishes were used within one week of PLL-g-PEG coating. For uniaxial compression of cells on PEG-coated dishes, we used the indentation mode of the Piuma control software (Optics11 Life), which allowed us to control the strain rate at which we deformed the sample. During measurements, the elapsed time, piezo displacement, and measured force were recorded at a time resolution of 1000 Hz. To address concerns that keeping our fibroblasts in suspension would affect cell viability, we measured the live-to-dead ratio of the cells and found no decrease over a period of 2 hours (the maximum length of an experiment) (Supplementary Figure S4). We calculated the contact area between the cell and the plates from epifluorescence microscopy images of the cytoplasm labeled with CellTracker orange (Fisher Scientific), where we thresholded each image using Otsu’s method in the Python library scikit image ^68^ and counted the number of pixels in the thresholded image. The applied stress was thus calculated as the measured force divided by the contact area. We calculated the uniaxial strain strain by approximating each cell as a sphere, and dividing the length of compression by the cell diameter. All measurements were made at 37°C, maintained by a microscope stage heater (Tokai Hit). For our polyacrylamide (PAA) benchmarking experiments, microgels with radii between 5 and 50 µm, ^69^ were immersed in MilliQ water for measurement. We selected beads with a diameter approximately equal to that of our cells (∼18µm). We applied the exact same experimental protocols to our PAA microgels as for cells to test our setup. After each experimental session, the cantilever was cleaned with 2-Propanol, rinsed with MilliQ water and carefully dried with a Kimtech tissue to remove organic waste.

### Dynamic mechanical analysis measurements

To measure the viscoelastic response of cells in the small deformation regime, we sinusoidally compressed them with the wedged microcantilever. First, the microcantilever was lowered towards the cell’s top surface, until the cell was compressed by 1 µm, corresponding to an approximate 5 % strain determined from the average cell size. After a waiting time of 20 seconds to allow for initial viscoelastic relaxation, cells were subjected to 200 nm oscillations (the minimum amplitude to obtain an acceptable signal-to-noise ratio) at 5 frequencies logarithmically spaced between 0.1 Hz and 1 Hz. From the applied strain and measured stress as a function of frequency, *f* , the complex viscoelastic modulus, *E*^∗^(*f* ), was calculated as

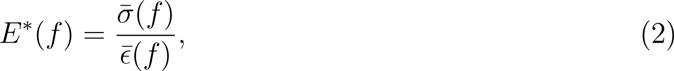

where *σ̄* and *ɛ̄* are the Fourier transforms of the time-dependent stress and strain, respectively. This equation tells us the frequency dependence of the cell mechanical response. We applied Equation 2 at each measured oscillation frequency by taking the fast Fourier transform (FFT) of the signal in Python, using the FFT function in Python’s numpy library. *E*^∗^(*f* ) data were decomposed into real (in phase) and imaginary (out of phase) parts, *E*^∗^(*f* ) = *E*^′^(*f* )+*iE*^′′^(*f* ), where *E*^′^(*f* ) and *E*^′′^(*f* ) are the compressive storage and loss moduli, respectively. *E*^′^(*f* ) corresponds to the elastic modulus of the cell (i.e., the energy stored per cycle of oscillation), whereas *E*^′′^(*f* ) corresponds to the viscous component (i.e., the energy dissipated per cycle of oscillation). We further decomposed Equation 1, the structural damping model, into real and imaginary parts:

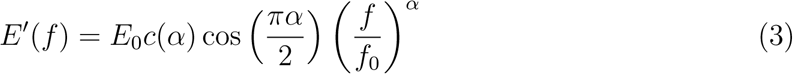

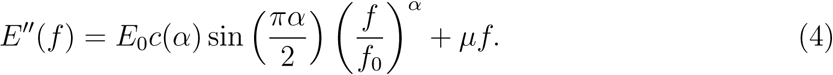

Using the curve fit function from Python’s SciPy^70^ library, we simultaneously fit Equations 3 and 4 to our elastic and viscous moduli, which yielded shared parameters for *E*_0_, *α* and *µ*.

### Strain-controlled compressive ramps

The same wedged microcantilever as used for dynamic oscillatory measurements was lowered towards a cell’s top surface in steps of 1 µm until cantilever deflection was detected, indicating contact with the cell. We then allowed the cell to relax for 30 seconds before beginning the strain ramp measurement. We chose to start the measurement from this contact position as we often noticed a small amount of slip upon initial contact with the cantilever. The cantilever was then gradually lowered, controlling the speed of cell compression at 2 µm/s unless specified otherwise (for rate dependence, see Supplementary Figure S1), until a strain of approximately 40% was reached. We checked by concurrent imaging that the cell reversibly changed shape (Supplementary Video 1). Above strains of 50%, we observed cell death in a large proportion of cases, as indicated by bleb formation visible in wide field microscopy (Supplementary Video 2). The cantilever was subsequently retracted at the same rate. The force acting on the cantilever was recorded during the entire loading-unloading cycle. To calculate cell stiffness, *K*, stress-strain data were numerically differentiated using the Python numpy library’s gradient function. ^71^ To determine the onset stress for stress-stiffening, we performed a piecewise linear fit to the stress/stiffness curve for each individual cell, where we fitted a horizontal line and a linear equation of the form *y* = *mx* + *c* with the Python scipy library, optimizing the onset point by maximizing the combined *R*^2^ values of both fits.

### Live cell imaging with confocal microscopy

Confocal microscopy to determine the localization and arrangement of the actin and vimentin cytoskeletal networks and the nucleus in cells was performed on a Leica Stellaris 8 LSCM utilising a 63x glycerol immersion objective (HC PL APO, WD 0.3 mm, NA 1.2), equipped with a white light laser operated at 488 nm (vimentin imaging) and 640 nm (actin imaging) and with a 405 nm diode laser (nucleus imaging). Detection was performed with the HyDS1 detector (440–480 nm, nucleus imaging), the HyDS2 detector (504-590 nm, vimentin imaging) and the HyDX3 detector (658-809 nm, actin imaging) operated in counting mode. To label vimentin, four days prior to imaging experiments, cells were transfected with vimentin-GFP^72^ using the Neon Transfection System (Thermo Fisher Scientific) with a single pulse width of 30 ms and a pulse voltage of 1350 V, according to the manufacturer’s instructions. We found that cells required four days to express vimentin at stable levels. For actin labeling, on the day of experiments, cell culture medium was replaced with fresh medium containing SiR actin (Tebu Bio, SC001) (1:1000) along with the pump inhibitor verampamil (Tebu Bio, SC001, part of SiR actin kit) (1:1000). Cells were incubated with the staining solution for 1 hour, before replacement with fresh medium, containing Hoechst 33342 (Thermo Fisher Scientific, H3570) (1:10,000) to label the nucleus. To mimic the conditions of our cell rheology experiments, cells were trapped between two coverslips coated with PLL-g-PEG, kept apart with 10 µm silica particles (Sigma, 904368) (see Supplementary Figure S5 for schematic). 3D confocal z-stacks were acquired with 256 x 256 pixel xy-slices separated by 0.33 µm at a scan speed of 400 Hz. The orthogonal projection tool in FIJI^73^ was used to generate side projections of cells. To generate ‘average cells’, a maximum projection was generated from a 2 µm section centered at the cell equator. Multiple-channel stacks (containing projections of the actin, vimentin and the nucleus) were aligned using HyperStackReg plugin, ^74^ which registered cells on top of each other, correcting for differences in cell size. In FIJI, we then made average intensity projections to generate an average cell. Finally, the radial profile plot tool in FIJI was used to calculate the normalized intensities around concentric circles from the center of the image. Images were collected from at least 2 independent experiments for each condition.

### Removal of nuclei from cells

We developed a protocol for enucleating the cells following a well-established protocol. ^47^ Cells were seeded on 22 mm diameter plastic coverslips and grown to 90% confluency in standard culture medium. Next, the cells were treated with cytochalasin D at a concentration of 1 µM to promote cell membrane flexibility and facilitate enucleation under centrifugation. After exposure to cytochalasin D for 30 minutes, the cells were placed in centrifuge tubes (344058 Beckman Coulter), on top of custom-made Teflon inserts (for schematics, see Supplementary Figure S6) facing downwards and centrifuged at 1000×g for 1 hour to remove the nuclei. Enucleated cells were transferred to fresh culture dishes and maintained in the same culture conditions to observe post-enucleation viability and behavior. Although we observed some enucleated cells survived up to 3 days post-enucleation, cells were used in experiments within 24 hours. Enucleation efficiency was checked by Hoechst staining before and after the procedure.

### Statistics and code availability

All mechanical experiments shown were repeated on at least three different days and imaging experiments were repeated twice. P-values shown to determine statistical significance were calculated using the paired Student’s t-test. Measured values in the main text are stated as the mean ± the standard error on the mean. Computer codes described in the Methods section are available on request.

## Acknowledgement

We gratefully thank J. Eriksson (University of Turku) for generously supplying the cells used in this study and R. Rodriguez and T. Schmidt (Leiden University) for providing the hydrogel beads. We thank M. Mavrakis (Institut Fresnel) for donating the vimentin-GFP plasmid used for live cell imaging experiments. We thank D. de Roos (TU Delft) for manufacturing the centrifuge tube inserts used for cell enucleation and N. van Vliet for help with electroporation. G.H.K. gratefully acknowledges funding from the NWO Talent Programme which is financed by the Dutch Research Council (project number VI.C.182.004).

P.E.B. gratefully acknowledges funding from the European Research Council (ERC) under the European Union’s Horizon 2020 research and innovation program (grant agreement no. 819424). F.C.M. was supported in part by the National Science Foundation Division of Materials Research (Grant No. DMR-2224030) and the Center for Theoretical Biological Physics (Grant No. PHY-2019745)

## Supporting Information Available

- Supplementary video 1.mov: bright field time lapse movie showing reversible loading and unloading of a cell to a compressive strain of approximately 0.45. Scale bars 10µm.
- Supplementary video 2.mov: bright field time lapse movie showing how excessive loading of a cell (above compressive strains of 0.5) leads to cell bleb formation. Scale bars 10µm.
- Supplementary information.pdf: contains the supplementary figures referred to in the main text.

