## Supplementary figures for "Actin and vimentin jointly control cell viscoelasticity and compression stiffening"

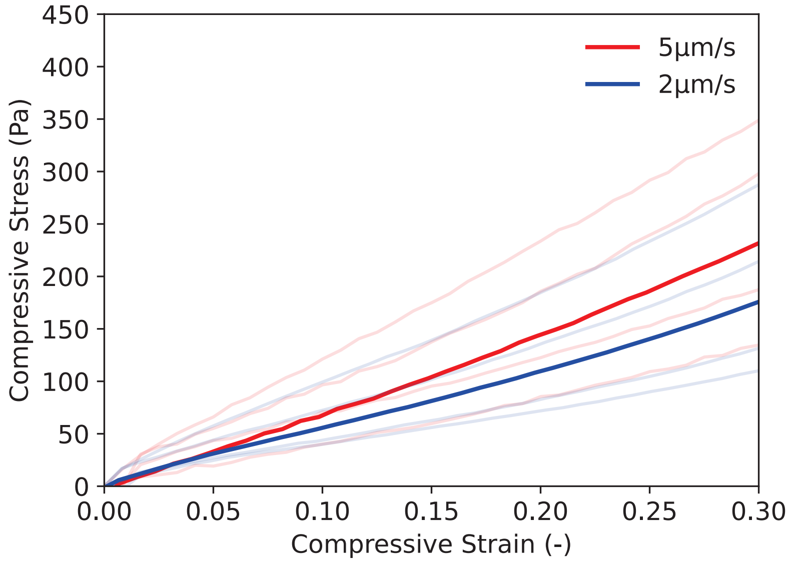


Figure 1: Experiments to determine the rate dependence of strain ramp tests. Compressing cells at a faster rate results in an apparently stiffer cell, consistent with general viscoelastic material properties. Wild type MEF cells were used for this experiment. Solid lines represent the average of all measurements and semi-transparent lines are the individual curves. N = 4 for both compression rates.

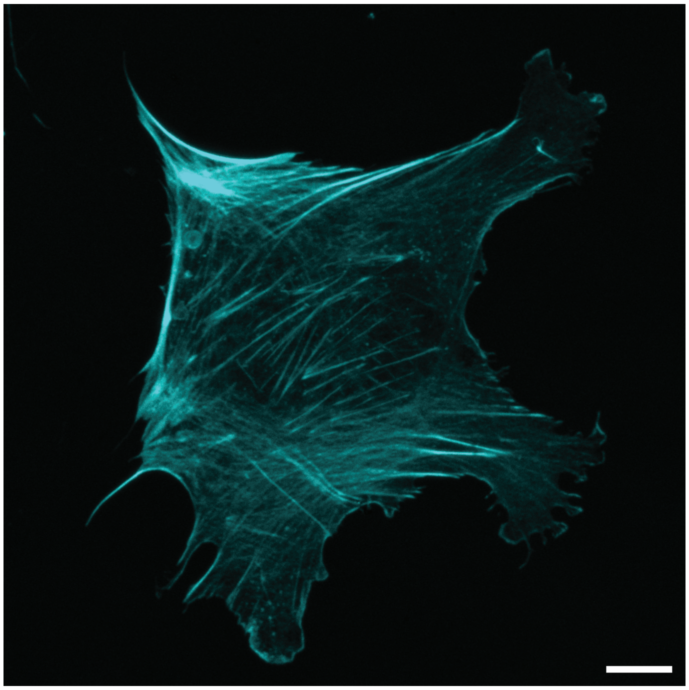

Figure 2: Representative maximum projection image (from a confocal z-stack with 32 slices separated by 0.33 µm) of a wild type mouse embryonic fibroblast adhered to fibronectin-coated glass and stained with Alexa-phalloidin to label F-actin. Adherent cells display pronounced stress fibres, in contrast to suspended cells (see images in main text). Scale bar 10 µm.

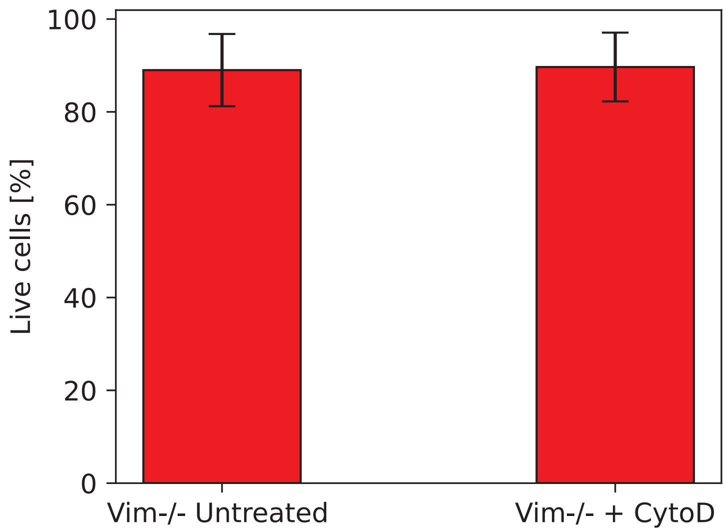


Figure 3: Live/dead comparison to test cell viability for vim -/- cells treated with cytochalasin D to depolymerize actin. Wild type mouse embryonic fibroblasts were stained with trypan blue to determine cell viability. Cell viability does not decrease after treatment with cytochalasin D. N=3. Error bars show the standard deviation.

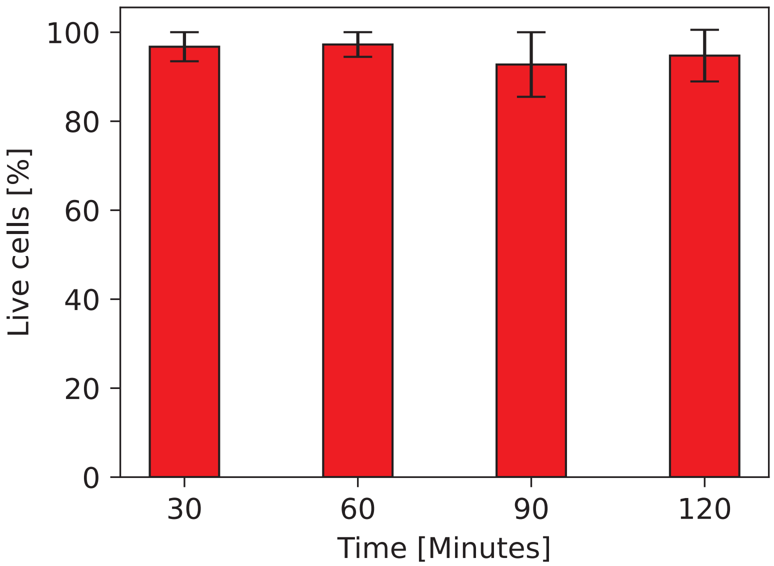


Figure 4: Live/dead comparison to test cell viability for the duration of a cell compression experiment. Wild type mouse embryonic fibroblasts were stained with trypan blue to determine cell viability. Cells were counted every 30 minutes up to 120 minutes (the maximum time cells were kept on the setup). Cell viability does not decrease during the course of an experiment. N=4. Error bars show the standard deviation.

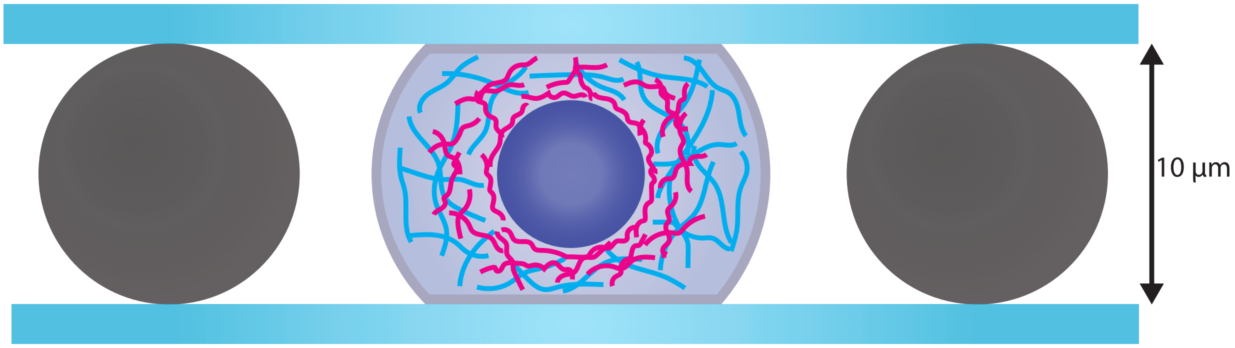


Figure 5: Schematic of the sample chamber used to acquire confocal images of single fibroblasts between two parallel glass coverslips to mimic the compression experiment conditions. Coverslips were coated with PLL-g-PEG to prevent cell adhesion, as in the compression experiments. Cells were trapped between coverslips separated by 10 μm beads.

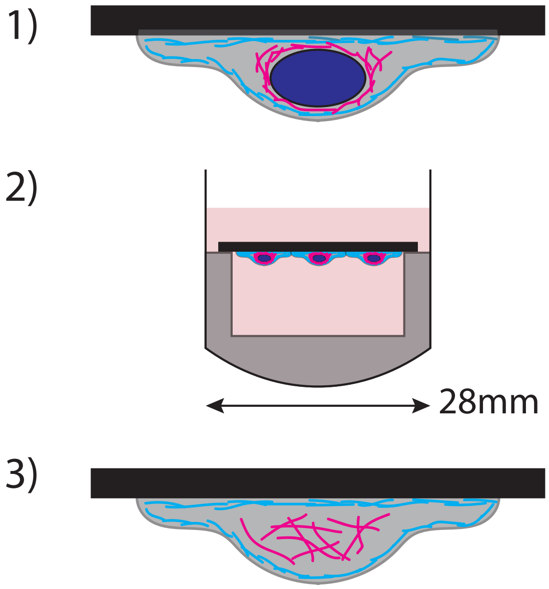


Figure 6: Schematic of the apparatus used to enucleate cells. 1) Cells are grown on fibronectin-coated plastic coverslips. 2) Cells are gently centrifuged faced down in a centrifuge tube with a teflon insert. 3) Enucleated cells remain adhered to the coverslip.

### Supplementary Methods

#### Actin staining for adherent cells

35mm glass bottom coverlips (Ibidi, 81158) were incubated with 20 µg/mL bovine fibronectin (Thermo Fisher Scientific, 33010018) for 1 hour at room temperature. Cells were subsequently seeded into dishes and allowed to adhere overnight. The next day, cells were fixed with 4% paraformaldehyde (PFA) (Thermo Fisher Scientific, J19943.K2) in phosphate-buffered saline (PBS) for 15 minutes at room temperature. Fixed cells were washed three times with PBS and then permeabilized using 0.1% Triton X-100 (Sigma, X100-5ML) in PBS for 10 minutes. After an additional three washes with PBS, cells were incubated with Alexa Fluor® 568-phalloidin (Thermo Fisher Scientific, A12380) diluted (1:1000) in PBS containing 1% bovine serum albumin (BSA) (Thermo Fisher Scientific, 15260037) for 30 minutes at room temperature, protected from light. Cells were then washed three times with PBS and subsequently imaged using the Leica Stellaris confocal microscope described in the main text.

#### Live/dead staining

For experiments showing cell viability throughout the length of an experiment (Fig S4), cells were harvested from the compression setup at 30-minute intervals. For experiments testing cell viability of vim -/- cells treated with cytochalsin D, cells were incubated in $CO_2$ -independent medium supplemented with 1 µM Cytochalasin D for 20 minutes. Cells were counted using a Countess Cell Counter (Thermo Fisher Scientific). For live/dead assessment, 10 µL of cell suspension was mixed with 10 µL of 0.4% trypan blue solution (Thermo Fisher Scientific, 15250061). Dead cells, which take up the trypan blue dye and appear dark or black under the microscope, were distinguished from live cells, which exclude the dye. The ratio of live-to-dead cells was recorded for each time point.
